# Odor cueing of declarative memories during sleep enhances coordinated spindles and slow oscillations

**DOI:** 10.1101/2023.02.10.527339

**Authors:** Andrea Sánchez-Corzo, David M Baum, Martín Irani, Svenja Hinrichs, Renate Reisenegger, Grace A Whitaker, Jan Born, Jens G Klinzing, Ranganatha Sitaram

## Abstract

Long-term memories are formed by repeated reactivation of newly encoded information during sleep. This process can be enhanced by using memory-associated reminder cues like sounds and odors. While auditory cueing has been researched extensively, few electrophysiological studies have exploited the various benefits of olfactory cueing. We used high-density electroencephalography in an odor-cueing paradigm that was designed to isolate the neural responses specific to the cueing of declarative memories. We show widespread cueing-induced increases in the duration and rate of sleep spindles. Higher spindle rates were most prominent over centro-parietal areas and largely overlapping with a concurrent increase in the amplitude of slow oscillations (SOs). Interestingly, greater SO amplitudes were linked to a higher likelihood of coupling a spindle and coupled spindles expressed during cueing were more numerous in particular around SO up states. We thus identify temporally and spatially coordinated enhancements to sleep spindles and slow oscillations as a candidate mechanism behind the benefits of odor cueing. Our results further demonstrate the feasibility of studying neural activity patterns related to memory processing using olfactory cueing during sleep.

**Statement of Significance:** Memory cueing during sleep allows insights into memory consolidation. This study is the first to investigate olfactory cueing-induced declarative memory processing using high-density EEG, while robustly controlling for critical confounding factors. The use of odors as cues, instead of more common auditory stimuli, further minimizes possible distortions due to sensory-evoked potentials. We demonstrate intricate changes in brain activity in response to cueing, such as the patterns of sleep spindles, slow oscillations, and their spatiotemporal coupling. We provide evidence that the enhancement of slow oscillation amplitudes, together with associated increases in sleep spindle rates, could be the key mechanism behind cueing-related memory benefits. We moreover show that prior findings obtained using auditory cueing are not the mere result of tone-evoked responses but might be genuine signatures of memory processing.

## Introduction

Recently encoded information must undergo active consolidation during sleep to be converted into long-term memories^1–4^. Reactivation results in a restructuring and strengthening of new memories, a process called memory consolidation. One way to probe this process is a technique called cued memory reactivation, in which sensory reminder cues, like odors^5, 6^ or sounds^7–9^ are presented during memory acquisition to associate them with the learned content. If the cues are presented again during subsequent sleep, this directs the process of consolidation towards the associated memories and improves their recall the next day^10^.

Auditory cueing is often considered more suitable for neuroscientific investigations^10^. Sounds more readily allow for high numbers of different cues within a single experimental night, they are independent of breathing, have clearly defined on- and offsets, and evoke stereotypical, time-locked neuronal responses that can be precisely analyzed. Cueing with odors in turn has other advantages. By bypassing thalamic relays, olfactory inputs reach the neocortex and hippocampus more directly than other modalities^11^. Furthermore, odors tend not to disturb sleep^12^ and lack of obvious evoked potentials eliminating an artificially induced factor that might confound the results. Sleep oscillations like slow waves and spindles are known to be linked with cognitive functions, including the sleep-associated consolidation of memories^1, 4^. While these oscillations are easily evoked using sounds, only few studies have demonstrated effects in these patterns in response to odor cueing^13–17^. Cueing effects on oscillatory patterns likely depend on the precise task to be memorized, cueing modality, and analysis technique. This is reflected in a spread of the demonstrated effects across various brain areas, frequency bands, and time delays relative to the cue, as well as some failed attempts to find robust associations between neural responses and memory improvements.

In this study, we aimed at closing this gap in our knowledge by carefully characterizing changes in neural activity in response to odor cueing of declarative memories. We focused on the main candidates for mediating beneficial effects of cueing, i.e., spindles, slow oscillations (SOs), and their interaction. For this purpose, we contrasted responses to two different odors. Before sleep, each odor was associated either with memories encoded during a declarative object-location memory task or with a procedural random finger-tapping task. The latter task did not contain any learnable sequences and was designed to, at most, induce sensory-motor learning. We conducted high-density electroencephalographic (EEG) recordings during subsequent sleep, while in each of two nights, either one of the two odors was presented. We compared neuronal responses to the two odors with the goal to unravel neuronal activity specific to the cueing of declarative memories. Memory performance was comparable after both nights, independently of the odor presented during sleep. Nevertheless, we found centro-parietal increases in the rate of sleep spindles and amplitude of SOs as an early response to declarative memory cueing. A follow-up analysis showed that higher SO amplitude was linked to more frequent SO-spindle coupling, pointing towards a mechanistic link of these two phenomena. Moreover, the duration of spindles increased over frontal areas. During SOs, spindle occurrence increased specifically around depolarized up states and their preferred phase was shifted slightly forward along the SO phase cycle. In summary, our results point towards an essential role of spindles, SOs and their spatiotemporal coupling not only in spontaneous^18, 19^ but also externally induced memory reprocessing. We further demonstrate the feasibility of investigating declarative memory-specific neural responses to odor cues^18, 19^.

## Methods

### Participants

Twenty-three young adults between 19 and 25 years of age (12 females; 22 ± 2.0 years, mean ± SD) participated in the study. Exclusion criteria were smoking, a diagnosed psychiatric disorder, ongoing medication, shift work, and a body-mass index (BMI) higher than 25. Participants were required to have a normal sleep-wake rhythm (i.e., going to bed between 10:00 p.m. and 1:00 a.m. and getting up between 6:00 a.m. and 9:00 a.m.). Participants were instructed to abstain from alcohol, naps, excessive exercise, extreme stress, as well as caffeine after 2 p.m. on the days of the experiment. Additional 29 participants were recruited but did not complete the study due to termination of the experiment for personal reasons or schedule incompatibility (n = 4), sleepless adaptation night (n = 3), illness (n = 3), insufficient learning performance (see below; n = 2), exceeding the maximum time to fall asleep (60 min; n = 1), insufficient deep sleep (< 20 min within 90 min of sleep; n = 14), or technical problems (n = 2). Participants received a compensation of 170 € for completing all experimental nights. The study was approved by the ethics committee of the medical faculty of the University Tübingen. All subjects gave their written informed consent.

### Experimental Design

The study consisted of an adaptation night and two experimental nights. The adaptation night allowed familiarization with sleeping with the high-density EEG system and face masks. Participants did not perform any memory tasks and were not presented with odor during that night. The adaptation night was performed 1-14 days before the first experimental night, and the second experimental night 14-28 days after the first. **Figure 1** shows a scheme of the experimental design.

**Figure 1:**
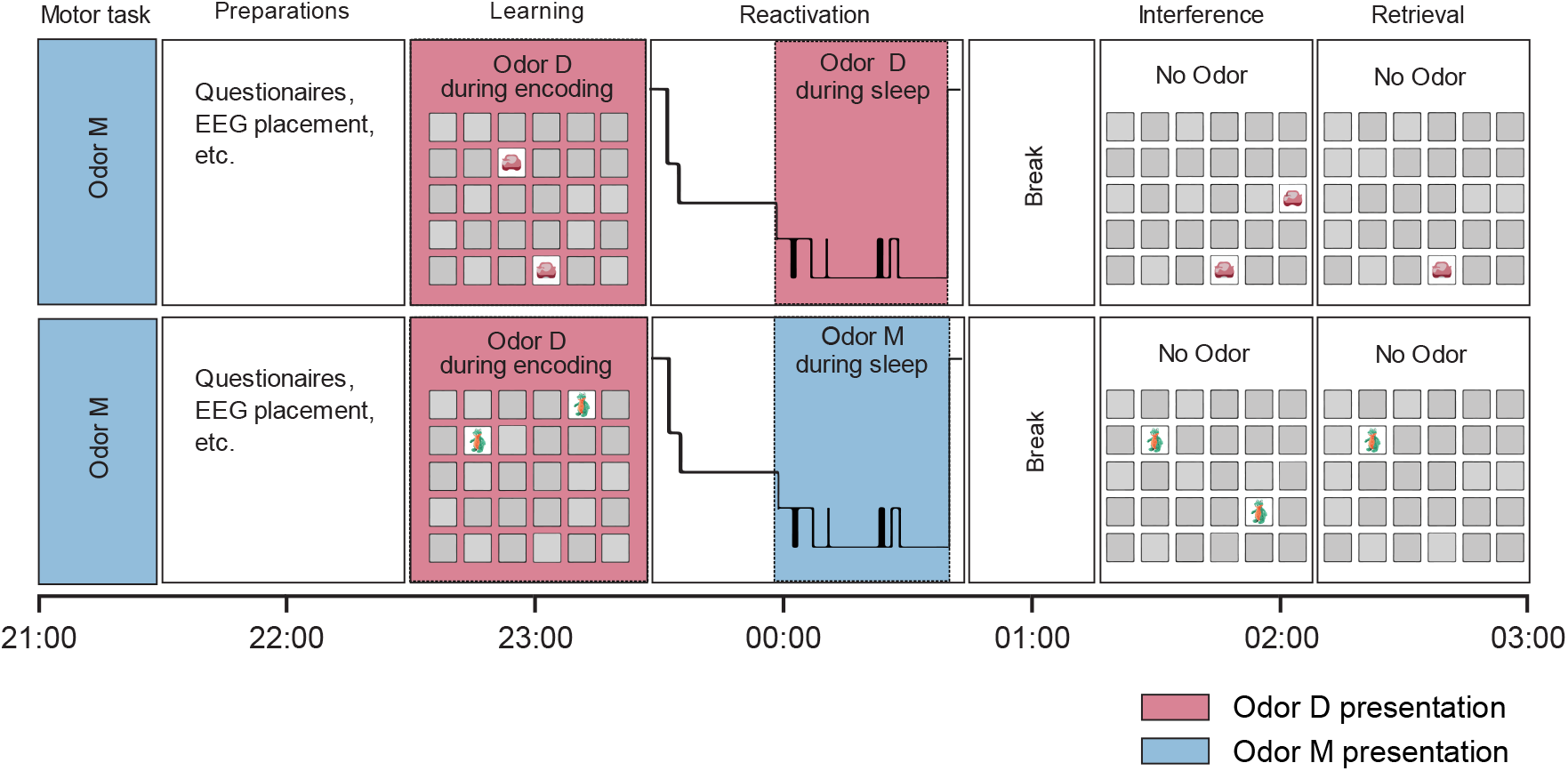
Experimental Paradigm. Participants completed two experimental nights in which they first performed a procedural Random Reaction Time Task in the presence of Odor M. After a break, participants learned a declarative memory task while administered with a different Odor D. Subjects went to bed and slept while their brain activity was recorded using high-density EEG. Once participants reached deep sleep, they received odor stimulation, either with Odor D or Odor M, depending on the condition (the order of which was counterbalanced across participants). Subjects were woken up and performed an interference task, after which retrieval performance on the original task was tested.

During experimental nights, participants first (at around 7:30 p.m.) performed a finger-tapping task with no learnable sequences (Random Reaction Time Task, RRTT). During this task, they were stimulated via a face mask with an odor defined as the “Motor task-associated odor” (“Odor M”). Two hours after the RRTT, participants learned the location of card pairs in a declarative memory task (see section ‘Declarative memory task’) while being stimulated with an odor defined as the “Declarative task-associated odor” (“Odor D”). During the learning period and after all cards had been presented twice, subjects completed the encoding of the image locations by performing an immediate recall test. The test was repeated up to 5 times until the participants answered more than 60 % of the cards correctly. The percentage of correct responses in the last run represents the accuracy of learned paired locations before going to sleep. If participants did not reach 60 % performance after the last run, the experiment was ended, and their data discarded (see ‘Participants’ section).

Upon reaching slow-wave sleep (SWS), either Odor D (in Night D) or Odor M (in Night M; balanced crossover design) was presented for windows of 15 s. Odor presentation was alternated with an odorless vehicle with 15-s breaks in between (resulting in a sequence of 15 s odor, 15 s break, 15 s vehicle, 15 s break, and so on). Stimulation was stopped when participants shifted from slow wave sleep (SWS) to another sleep stage or woke up. Once participants had slept for a maximum of 90 minutes and received between 40 and 80 stimulations (either odor or vehicle), subjects were woken up and were shown a movie for 30 min to allow shaking off sleep inertia. Subsequently, they learned an interference task (see below) without any odor presentation. This was followed by another 45-min movie break before testing the participant’s performance on the pre-sleep declarative memory task. Participants were tested for a single run. The experiment ended between 2:00 and 4:00 a.m. The subjects slept until the morning without EEG monitoring or further experimental procedures.

For analyses not reported here, participants also underwent an anatomical magnetic resonance imaging scan before the experiment and three 10-min resting-state recordings throughout each experimental night. Citral and Isobutyraldehyde (Merck Sigma Aldrich) were used for Odors D and M in a counterbalanced fashion.

### Random Reaction Time Task (RRTT)

Participants performed a motor task with visual cues (four different displayed keys) as instructions for pressing corresponding keys on a keyboard with predefined fingers (excluding the thumb). The task was similar to the classic "Serial Reaction Time Task" but without any predefined repeating patterns. This made it impossible to improve performance by learning any sequences of keys, which was meant to minimize the encoding of new memories other than sensory-motor learning. Incorrect keystrokes were signaled to the participant. The test was performed in two blocks of five minutes each with a short break in between. Throughout the task, the motor task-associated odor (Odor M) was presented to the participant in an intermittent fashion (15 s on, 15 s off). One goal of the RRTT was to expose the participants to Odor M during an engaging task and thereby make it comparable to Odor D in terms of pre-sleep familiarity. Task length and odor stimulation were chosen to structurally mimic the declarative memory task.

### Declarative memory task

Participants also performed a two-dimensional object-location memory task used in previous studies^5, 6^. This task is an adaptation of the social game "Concentration" or "Memory". A board with 30 gray squares in a 5 x 6 matrix represented 15 face-down pairs of cards. Cards showed everyday objects and animals in color. For the two nights, two different sets of cards and card locations were used. During *learning*, card pairs were revealed one by one, such that the first card was shown for 1 s after which also the second card was revealed for another 3 s. Next, both cards were turned face down and the next card pair was revealed. This continued until all card pairs were presented twice. For a subsequent *immediate recall test*, subjects were presented with the first card of a pair and were asked to indicate the location of the second card using the mouse pointer. Correct responses triggered a green tick mark, which appeared for 1 s at the location of the selected card. Incorrect responses triggered a red cross and the correct card location was revealed for 3 s. Immediate recall was repeated until the participants showed a performance of at least 60 % correct responses (learning criterion) but not more often than 5 times. In a later *interference* version of the task, participants repeated the learning procedure using cards with the same images but the second card of each pair was placed at a different location^20, 21^. This AB-AC learning scheme (A, B, and C correspond to different localizations) was used to test the stability of the original memory against partly overlapping and thereby interfering information. During the interference task, the subjects wore the odor mask but no odor was applied. Finally, participants performed another single run of the recall test using the card locations of the original task version. No direct feedback was given during the final *recall*. Only at the very end, the overall performance was shown. Task performance was measured by the percentage of the number of correctly recalled card pair locations after sleep (recall test) relative to the number of correctly recalled pair locations before going to sleep (learn lest) as a percentage: Recall Test / Learn Test * 100.

### Data acquisition

EEG data were acquired using a 129-electrode high-density EEG system (Electrical Geodesics Inc., Eugene, United States) with electrodes placed according to an adapted 10-10 system (HydroCel Geodesic Sensor Net). Data were recorded at a sampling rate of 1000 Hz and online-referenced to the vertex electrode. The location of each electrode in relation to the head was registered using optical tracking (Localite TMS Navigator, Localite GmbH, Bonn, Germany). For this purpose, structural T1-weighted magnetic resonance images were acquired, which allowed reconstructing each participant’s individual head shape (1 mm^3^ voxel size; MAGNETOM Prisma^fit^ scanner, Siemens Healthcare GmbH, Erlangen, Germany). In the beginning of the first experimental night, three optical references (small reflective tracking balls with defined relative positions, provided by the manufacturer) were attached to the participant’s head. This was done by fixing them to an individually molded martial arts mouth guard, which allowed a precisely reproducible connection to the head. The reference balls, as well as a large number of points on the scalp were localized in space using an optically trackable pointing device. Reference balls and scalp locations were then coregistered with the surface of the reconstructed head model. Subsequently, the positions of all EEG electrodes were registered and added to the resulting common coordinate system. In the beginning of the second experimental night, mouth guard and attached tracking balls were placed again, correspondence with the participant’s head surface was validated, and all EEG electrodes were placed at their respective location in the first experimental night. This procedure ensured that each electrode was placed over the same brain area in both sessions.

### Sleep scoring and sleep architecture

Data were scored according to the Rechtschaffen & Kales (1968)^22^ scoring criteria based on channels equivalent to C3/C4, referenced to the averaged mastoid channels, as well as bipolar electrooculography (EOG) and electromyography (EMG). To aid sleep scoring, data were bandpass-filtered between 0.3-35 Hz and downsampled to 200 Hz. Each 30-s epoch was then labeled as Awake (W), Stage 1-4 (S1-S4), Rapid Eye Movement sleep (REM, R), or Movement Time (MT).

In order to evaluate the impact of cueing on sleep architecture, we compared the time spent in each sleep stage as well as sleep onset latency, time awake after sleep onset, number of arousals and sleep efficiency defined as the percentage of sleep after lights out.

### Pre-processing of EEG recordings

EEG data were preprocessed using the FieldTrip toolbox^23^. Datasets were first segmented into epochs of -5 to 15 s around stimulus onset. Epochs with a stimulation duration of less than 15 s were discarded (2.46 ± 2.54 %, mean ± SD). Subsequently, data were lowpass filtered at 30 Hz using a finite impulse response (FIR) filter. An additional median filter was applied with an order of 30 to suppress occasional non-sinusoidal technical artifacts. Electrodes located on the face or neck were discarded from further processing. Noisy EEG channels were visually identified and interpolated using a distance-based weighted average of the neighboring channels (proportion of interpolated channels: 1.14 ± 2.72 %, mean ± SD). All data were visually inspected for artifacts. Identified artifacts led to rejection of the entire trial in case several channels were affected (1.67 ± 5.32 %, mean ± SD), while in the case of short, channel-specific artifacts, trial-specific channel interpolation was performed (3.28 ± 7.52 %, mean ± SD). For all subsequent analyses, data were re-referenced to the averaged signal from both mastoid electrodes. In datasets where mastoid channels were identified as noisy, the channel closest to the affected one was used as a replacement. Finally, data were downsampled to 250 Hz.

### Event detection

Peak sleep spindle frequencies differ substantially from person to person. The peak frequency of spindles for each participant was identified individually using data from the central electrodes, where fast spindles are usually most pronounced. Irregular Resampling Auto-Spectral Analysis (IRASA)^24^ was performed to separate the fractal from the oscillatory component for all data recorded during NREM sleep. For each participant and recording night, the maximum power of the oscillatory component in the frequency band between 12 and 16 Hz was used as the spindle peak frequency (**Supplementary Figure 2A**). This procedure reliably resulted in fast spindle peaks (13.79 ± 0.38 Hz, mean ± SD). Secondary, slow spindle peaks were visible for some participants but were too unreliable across the entire sample to be used for further analysis.

Discrete sleep spindles were detected separately for each channel using a MATLAB custom script based on published methods^21^. The EEG signal was bandpass-filtered around the detected individual spindle peak frequency (± 1.5 Hz). The signal’s root-mean-square (RMS) was calculated using a sliding window of 0.2 s with a step size of one sample. Additional smoothing was performed with a sliding-window average of the same size. Time frames were considered spindle candidates if I) the RMS signal exceeded the value of the mean + 1.5 standard deviations (SDs) of the RMS signal of all combined stimulation epochs for a period between 0.5 and 3 s, and II) the RMS signal inside this period crossed a second amplitude threshold placed at mean + 2 SDs for at least one sample.

Individual slow oscillations (SOs) were detected automatically using a custom script based on spectral content and duration. The detection procedure was largely based on prior literature^21^. In detail, the EEG signal was bandpass-filtered to the slow-wave (SW) frequency range (0.5 - 4 Hz) using a zero-phase two-pass Butterworth filter with an order of 3. Candidates for SOs were identified by the presence of two consecutive negative-to-positive zero crossings, framing a negative and a positive half-wave, separated by the positive-to-negative zero-crossing inside this time window (**Supplementary Figure 3**). These candidates were further validated using three required characteristics: I) Negative half-waves needed to cross a threshold trough amplitude of 1.25 SDs from the average of the combined signal of analyzed trials; II) peak-to-peak amplitudes of candidates needed to cross a threshold of 1.25 SDs from the average peak-to-peak amplitudes of the combined signal of analyzed trials; and III) consecutive positive-to-negative zero-crossings needed to be separated by 0.8 to 2 seconds. From the resulting set of detected SOs, outliers with trough amplitudes beyond the median ± 3 SDs across all detected slow oscillations were discarded. The SO slope, as an established metric, was defined as the average of the slopes over samples between the negative peak and positive peak of the SO.

### Validation of detected events

The spindle and slow oscillation detection procedures were evaluated based on: I) The topographic distribution of the detection rate of events across the scalp (**Figure 2A** and **3A**); II) the average waveform and spectral composition of the raw signal (**Figure 2B** and **3B**) incl. where spindles reached their peak amplitude (**Figure 2C**); III) the occurrence rates of events across sleep stages (**Figure 2D** and **3C**); IV) the occurrence of detected spindles along detected slow oscillations (i.e., proportion of SOs having a nested spindle (**Figure 4B**) and phase-amplitude coupling (**Figure 4C** and **4D**); spindle time points were based on the event’s midpoint, half way between its on- and offset). Evaluations were done for all events during S2-S4.

**Figure 2:**
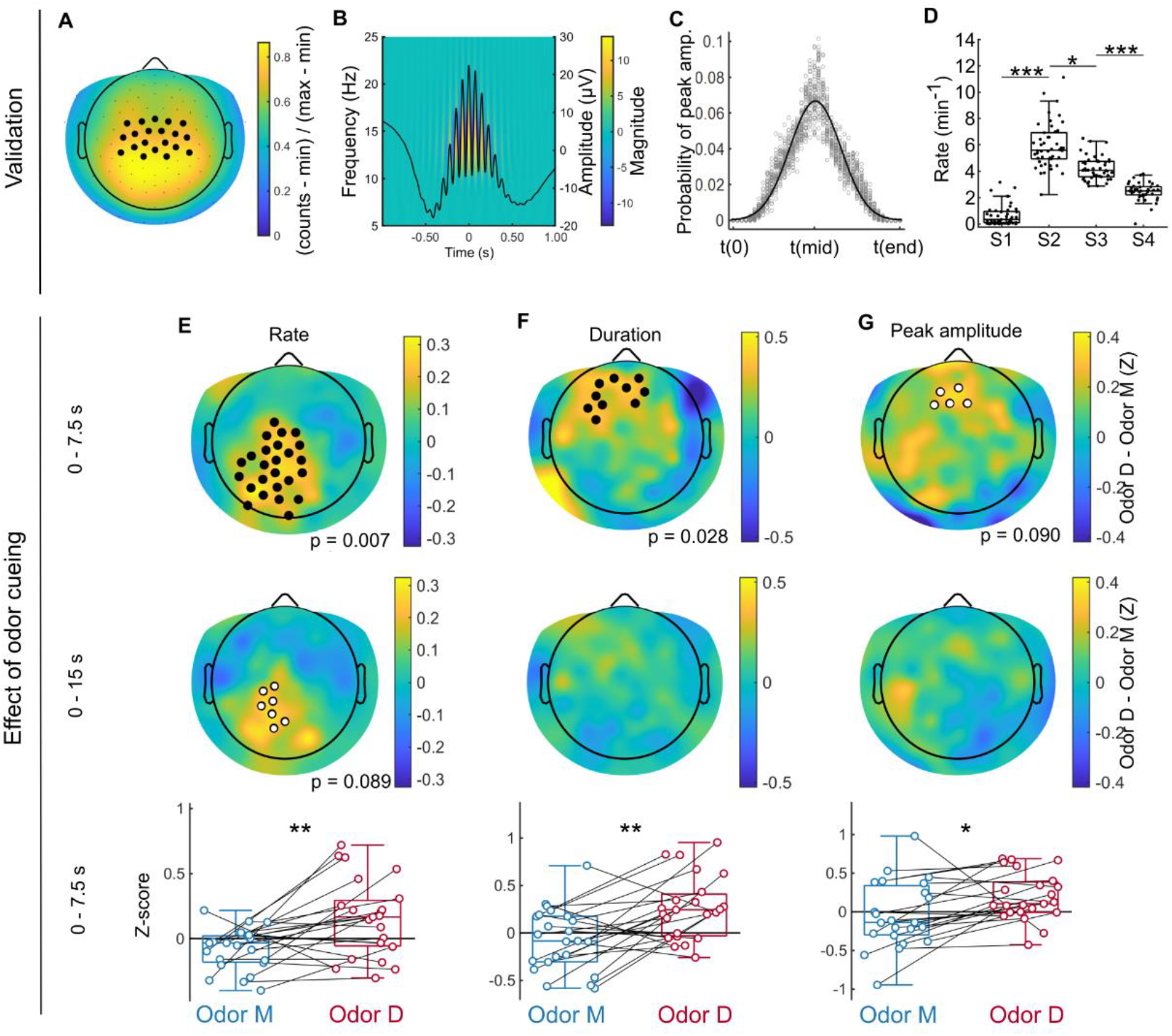
Odor cueing enhances several characteristics of fast sleep spindle activity. Detected sleep spindle events were validated based on established characteristics such as distribution of detection rate across scalp electrodes (**A**), average waveforms and time-frequency composition (**B**), location of spindle signal peaks (**C**; t(0), t(mid) and t(end) refer to onset, midpoint and offset times), as well as their detection rates across different sleep stages (**D**). Black circles in A represent electrodes belonging to the anatomically defined central cluster used for validation steps in **B**-**D**. Cueing with Odor D (red) compared to Odor M (blue) was associated in the first half of the stimulation period (0 - 7.5 s; upper panels) with significant increases in spindle rate (**E**; single cluster, see cluster-level p-value in panel) and duration (**F**), while peak amplitude did not differ significantly (**G**). These effects were not detectable when analyzing the entire stimulation window (0 - 15 s; **E**-**G** middle panels). Black circles over topoplots in panels E-G represent electrodes inside significant clusters (p < 0.05), white circles electrodes inside clusters with p-values < 0.10. Boxplots (**E**-**G** bottom panels) display changes between Odor D and M in 0 - 7.5 s averaged over the clusters shown in the top panels. ***, p < 0.001; **, p = 0.007 and 0.002 for **E** and **F**; *, p = 0.017 and 0.027 for **D** and **G**, respectively. Lines connect the same subjects across conditions.

**Figure 3:**
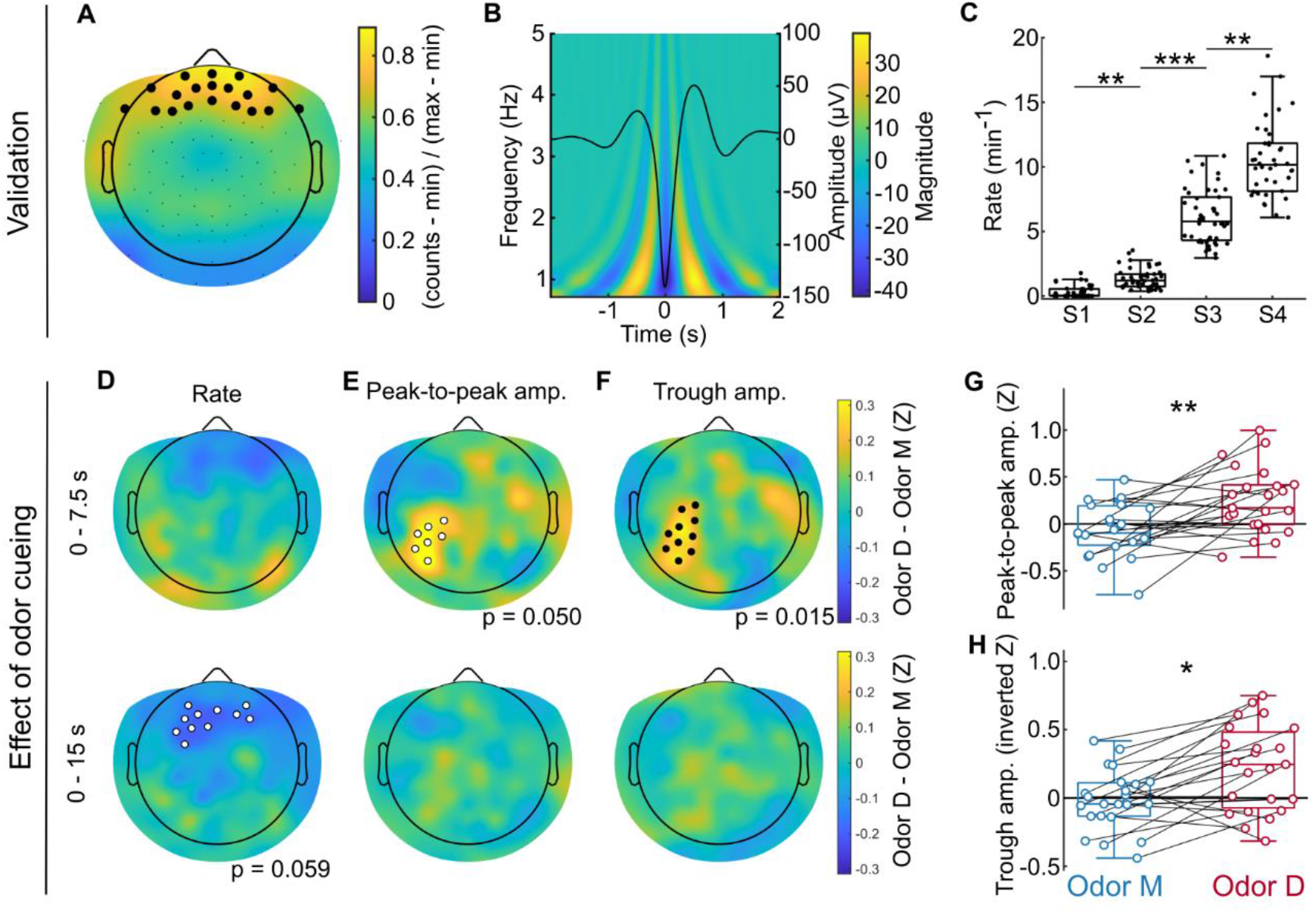
Declarative memory odor cueing increased the amplitude of centro-parietal slow oscillations. Detected slow oscillatory events were validated based on their canonical characteristics using their distribution across the scalp (**A**), their averaged waveforms and time-frequency composition (**B**), as well as detection rates across different sleep stages (**C**). Black circles in **A** represent electrodes belonging to the anatomically-defined frontal cluster used for validation steps in **B** and **C**. Presenting Odor D did not evoke changes in event detection rate (**D**) when compared to Odor M, but resulted in more negative SO peak-to-peak amplitudes (**E**) and trough amplitudes (**F**, Z-scores are inverted for visual clarity) specifically in the first half (0 - 7.5 s, top panel) of the stimulation window (but not in the whole window, 0 - 15 s, bottom panel). This effect is further illustrated by comparing averages across all electrodes inside the found cluster (**G** and **H**, data shown for 0 - 7.5 s, lines connect data from the same participant). *, p = 0.018; **, p = 0.004 (**G**), 0.003 (S1 vs S2) and 0.009 (S3 vs S4); ***, p < 0.001.

**Figure 4:**
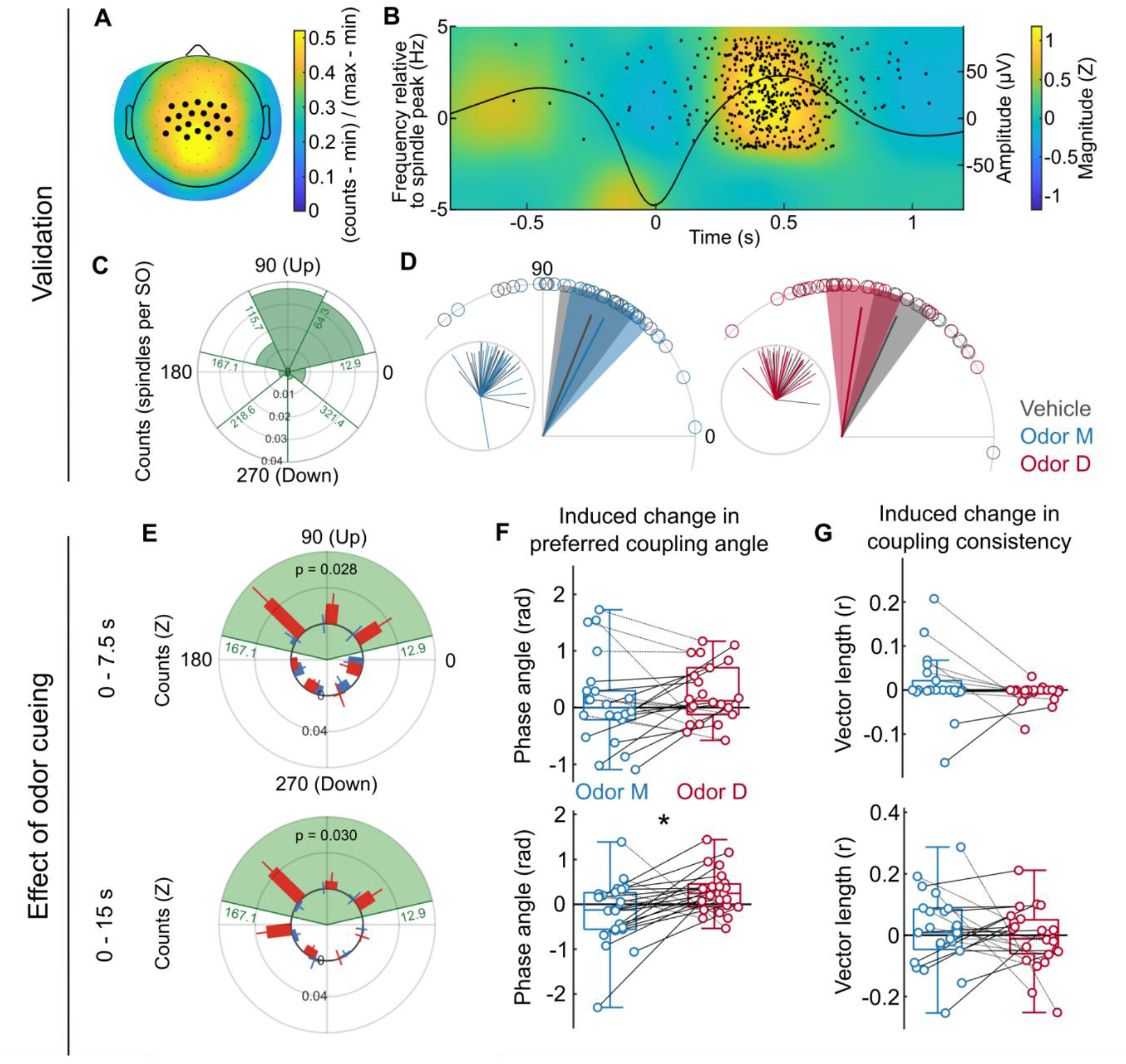
Declarative odor cueing induces sleep spindles specifically around slow oscillation up states. **A**, Most coupled events were detected in and around central electrodes (colors show spindle counts per slow oscillation, black circles show analyzed electrodes for all other panels of this figure). **B**, Time-frequency representation of spindle activity around slow oscillations. Shown is the power over central areas at frequencies ± 5 Hz centered around the average spindle peak frequency across subjects (13.80 Hz). Black waveform shows grand average slow oscillation. Black dots indicate spindle midpoints taken from the first half of each stimulation period in one night, pooled across all subjects (arbitrary vertical distribution). **C**, Spindles were mainly enhanced during SO up states. Green bars represent the average number of spindles in subjects for each phase bin across all detected SO events, participants, and nights (first half of stimulation periods; bin size: 51.4°; bin edges marked in green). **D**, Preferred phase angle (lines) and 95 % confidence intervals (areas) across participants. Data for each participant is shown as circles on the polar axis and lines in inlays. Line lengths represent coupling strengths. **E**, Changes in the number of spindles occurring during different SO phases (Z-scores ± SEM), separated by cueing condition. P-values are provided for the comparison of the averages across bins during the SO up state (between 12.9° and 167.1°, green background) which is also the phase range considered in **F** and **G**. **F** and **G** show values of preferred SO phase angles of coupled spindles and phase coupling consistency (vector length, r) as differences between odor and vehicle control periods. Upper panels in **E**, **F** and **G** show results for coupled events inside the first half of stimulation periods (0 - 7.5 s) in upper panels and for the whole stimulation periods (0 - 15 s) in lower panels. Lines connect data points from the same participant. *, p = 0.027.

### Event-coupling: spindles and SO

Slow oscillations tend to couple sleep spindles to their up phase. To analyze changes in this coupling, the signal was bandpass-filtered between 0.5 and 2 Hz and Hilbert transformed. Based on the resulting analytic representation of the signal, the phase was extracted at the midpoint of each detected sleep spindle (halfway between its on- and offset)^25^. Spindles were considered to occur along an SO when their midpoint was located between the SO onset and offset (positive-to-negative zero crossings). The relationship between SO amplitude and spindle coupling was assessed using SO amplitudes that were normalized by the mean and SD amplitude of all SOs occurring inside sham trials in the Odor D night. The resulting z scores were averaged across all SOs that nested a spindle or those that did not, as well as for all channels inside either of two analyzed channel clusters.

### Statistical analysis

If not otherwise noted, all statistics were performed using two-tailed tests and reported p values were corrected for multiple comparisons.

#### Event detection and features

Occurrences of spindles and SOs across sleep stages were analyzed by averaging their rates over channels within anatomically defined clusters and comparing them between sleep stages using the Kruskal-Wallis test. For analyzing differences in event characteristics between conditions, metrics obtained during odor stimulation were Z-scored using the mean and SD obtained across all vehicle trials of the same night. As a result, each odor trial was normalized by a common vehicle baseline. The two odor conditions were then compared using either a whole brain analysis or a confined subset of electrodes. The latter was done in cases where I) there was a clear hypothesis about the anatomical distribution of a potential effect (anatomically defined cluster) or II) a post-hoc analysis of an effect found in the whole-brain analysis was performed. For whole-brain analyses, channels with significant differences between conditions (two-tailed paired-samples t-tests; sample-level alpha 0.05) were grouped into connected clusters and cluster-level statistics were calculated by summing up the t-values within each cluster. Subsequently, the found clusters were subjected to multiple comparison correction by comparing them to a reference distribution obtained using a Monte Carlo permutation approach (cluster-based permutation analysis as implemented in Fieldtrip; maxsum method; cluster-level alpha 0.05; 100,000 permutations)^26^. For analyzing channel subsets, Wilcoxon’s rank sum tests were performed on the average value across all channels in that cluster.

#### Event coupling

Differences in the coupling of spindles to SOs were analyzed by comparing I) the number of coupled spindles, II) the phase consistency of this coupling, as well as III) shifts in the coupling phase. For all types of analysis, samples were normalized subject-wise by subtracting the average of all vehicle trials from the average of all odor trials of the same night. For coupling consistency analysis specifically, differences in the numbers of detected events and therefore phase estimates (potentially leading to systematic biases^27^) were controlled for by running the analysis on 10,000 randomly selected subsets of equal size and averaging the results. Samples were compared using channel cluster-based analysis as described in the previous section. Angular data obtained from the phase-coupling analysis were processed using the circ_stat toolbox^28^ and analyzed with the Watson-Williams multi-sample test. The amplitudes of SOs that coupled a sleep spindle and those SO that did not were compared across participants using the Wilcoxon signed-rank test.

#### Behavioral performance

Differences in memory task performances between odor conditions were evaluated using the non-parametric paired Wilcoxon rank sum test.

#### Samples

Data were considered outliers if their normalized values were more than 3 SDs above or below the median difference between all normalized Odor D and Odor M values. Such data points were removed before statistical analysis, together with their sample-specific counterpart of the contrasted condition. Outlier rejection was not performed for the analysis of coupled events due to the low number of trials.

## Results

### Odor cueing of declarative memories enhanced spindle activity

We analyzed the effects of odor cueing of declarative memories on the detection rate, duration, and amplitude of sleep spindles detected during NREM sleep. We found that, in the first half (0 - 7.5 s) of the stimulation period, declarative memory cueing increased the rate of sleep spindles over centro-parietal brain areas (Odor D vs Odor M, cluster-level p = 0.007, **Figure 2E**. For a time-resolved analysis, see **Supplementary Figure 4**), as well as their duration over frontal areas (p = 0.028, **Figure 2F**). These effects were weaker and not statistically significant when considering the whole stimulation period (0 - 15 s, lowest cluster-level p = 0.089 for spindle rate, **Figure 2E**). To further illustrate these changes and estimate their effect size, we conducted a follow-up analysis on averages across electrodes within found clusters (Odor D vs Odor M, spindle rates: 0.15 ± 0.06 vs -0.08 ± 0.03, average ± SEM, p = 0.007, Wilcoxon’s z = 2.702, **Figure 2E**; duration: 0.27 ± 0.07 vs -0.07 ± 0.0.07, p = 0.002, z = 3.098, **Figure 2F**; peak amplitudes: 0.20 ± 0.06 vs -0.04 ± 0.09, p = 0.027, z = 2.219, **Figure 2G**).

The process of detecting sleep spindles was assessed in a post-detection validation process illustrated in **Figure 2A-D** and the **Supplementary Figure 2**. As expected, I) detection rate of spindles was highest in centro-parietal electrodes (**Figure 3A**), II) the averaged signal showed a fusiform oscillatory activity in the sigma frequency band **(Figure 3B**), III) the maxima of the RMS signal preferably located around the event’s midpoint (**Figure 3C**) and IV) detection rates were highest in S2 and decreasing towards S4 (S1: 0.65 ± 0.12, S2: 5.93 ± 0.25, S3: 4.22 ± 0.13, S4: 2.47 ± 0.11, average ± SEM, **Figure 3D**).

### Declarative memory cueing enhances the amplitude of slow oscillations

After demonstrating that declarative memory odor cueing affected several characteristics of sleep spindles, we asked whether similar effects were apparent for slow oscillations. Indeed, declarative memory odor cueing resulted in an increased amplitude of slow oscillations over centro-parietal areas during the first half (0 - 7.5 s) of the stimulation period (Odor D vs Odor M: peak-to-peak amplitude, cluster-level p = 0.050, **Figure 3E**; trough amplitude, p = 0.015, **Figure 3F** with inverted Z scores for visual consistency between event features). Similar to the cueing effects on spindles, the effects on SOs were weaker and not statistically significant when analyzing the entire stimulation period (0 - 15 s). A follow-up comparison of the average across all electrodes in the found cluster shows the size of this effect for peak-to-peak amplitude (Odor D vs Odor M: -0.28 ± 0.07 vs -0.07 ± 0.06, average ± SEM, p = 0.004, Wilcoxon’s z = 2.856, **Figure 3G**) and trough amplitude (-0.22 ± 0.07 vs 0.02 ± 0.02, non-inverted z scores, p = 0.018, Wilcoxon’s z = 2.373, **Figure 3H**). Because the increases in spindle rate (**Figure 2E**) and SO amplitude (**Figure 3E and 3F**) occurred over similar areas, we asked whether these two phenomena might be related. Indeed, SO amplitude was generally linked to the likelihood of the occurrence of a coupled spindle. Independent of any cueing, SOs that co-occurred with a spindle tended to exhibit higher trough amplitudes compared to SOs that did not (Wilcoxon signed-rank test; central anatomical cluster shown in **Figure 2A**, p = 0.003; cluster of cueing-induced spindle rate increase, **Figure 2E**, p = 0.030).

The detection of SOs was evaluated similarly to detected spindles in a post-detection validation (**Figure 3A-3C**, **Supplementary Figure 3**): As expected, I) detection rate of SOs was highest in frontal electrodes (**Figure 3A**), II) the averaged signal showed SO-typical morphology with a strong negative peak in the negative halfwave (**Figure 3B**), and III) detection rates were steadily increasing from S1 towards S4 (S1: 0.30 ± 0.01, S2: 1.36 ± 0.02, S3: 6.13 ± 0.05, S4: 10.50 ± 0.07, average ± SEM, **Figure 3C**).

### Declarative memory cueing increases the number of spindles co-occurring with SOs

The cueing-induced changes in spindle rate and SO amplitude shown above both occurred over centro-parietal areas and were strongest in the first seconds after cueing onset. We therefore asked whether cueing further affected the more fine-grained temporal synchronization of these events. We analyzed the number of spindles occurring around detected SOs and found that this number was greatest over central areas (**Figure 4A**). We therefore performed subsequent analyses on averages across electrodes inside an anatomically defined central cluster. We found that slow oscillations and sleep spindles showed robust temporal synchronization, such that SOs tend to nest sleep spindles to their depolarized up state (**Figure 4B-D**). Of all SOs detected in central electrodes, 11.4 ± 0.4 % (average ± SEM across subjects) had spindles that matched our criteria throughout sleep stages S2 to S4. As expected, most SO-coupled spindles during trials occurred around the SO up state. To quantify this phenomenon, we divided the SO cycle into seven phase bins of 51.4° and assessed the number of spindles for each bin. 80.9 ± 1.73 % of those spindles that were broadly coupled to an SO (see Methods) had their midpoint during SO up states (i.e., occurred in one of the three phase bins spanning angles between 12.9° and 167.1°; **Figure 4C**). Importantly, while this up state coupling was present in both odor conditions (see **Figure 4D** for the average preferred coupling angle for each participant, separated by Odor D and M), declarative memory odor cueing further increased the number of spindles specifically around SO up states (between 12.9° and 167.1°). This was the case for the first half (Odor D vs Odor M: 0.032 ± 0.011 vs 0.003 ± 0.009, average ± SEM, p = 0.028, z = 2.195, **Figure 4E**, upper panel) and whole stimulation periods (Odor D vs Odor M: 0.025 ± 0.009 vs 0.001 ± 0.006, p = 0.030, z = 2.175, **Figure 4E**, lower panel) of the stimulation period. For phase bins outside the SO up state, spindle counts did not change significantly (first half of stimulation period, -0.012 ± 0.004 vs -0.012 ± 0.003, p = 0.878, z = 0.154; whole stimulation period: 0.005 ± 0.006 vs 0.000 ± 0.005, p = 0.279, z = 1.082).

Interestingly, cueing also resulted in a forward shift of the preferred phase angle at which spindles were coupled to the SOs. This effect was significant when considering the whole stimulation period (Odor D vs Odor M, 0.280 ± 0.101 vs –0.172 ± 0.150, average ± SEM, p = 0.027, F = 5.258, **Figure 4F** lower panel) but remained unchanged when only including events from its first half (0.247 ± 0.112 vs 0.107 ± 0.165, p = 0.362, F = 0.850, **Figure 4F** upper panel). The phase-specific spindle increase did not alter the coupling consistency (**Figure 4G**), probably because of the wide breadth and symmetry of this effect, respectively.

### No significant differences in memory performance, sleep architecture, or sleep quality

While electrophysiological features revealed changes when comparing the two stimulation conditions, there were no significant differences in memory performance (Night D vs Night M: 75.61 ± 5.38 vs 75.13 ± 4.22, average ± SEM (performance), p = 0.956, Wilcoxon’s z = 0.055, **Supplementary Figure 1**). Also, the distribution of sleep stages was comparable between conditions (Table 1). On average, subjects showed slightly more N1 in the Odor D night (Night D vs Night M: 6.61% vs 3.92%, uncorrected p = 0.008; see Table 1), with this small difference (0.28 min) being very unlikely to have any confounding effects on our results. Subjects tended to be in deeper sleep stages (S4, Night D vs Night M: 17.42 vs 22.59, average (%), uncorrected p = 0.052) and to spend less time awake after sleep onset (WASO, Night D vs Night M: 3.74 vs 2.35, average (%), uncorrected p = 0.059). Please note that all these tests are reported uncorrected for multiple comparisons to maximize statistical sensitivity.

**Table 1.** Sleep architecture across experimental conditions. Conditions did not differ in sleep macrostructure, incl. the proportion of sleep spent in each sleep stage. Arousals, Number of awakenings after sleep onset; REM, Rapid Eye Movement sleep; S1-4, Non-Rapid Eye Movement sleep stages 1 to 4; SE, Sleep efficiency (percentage of sleep after lights out); Sleep onset, Latency of first sleep score after lights out; SWS, Slow-wave sleep; TST, Total sleep time; WASO, Wake after sleep onset. Values are expressed as averages ± SEM. Only one significant difference emerged (S1, p = 0.008 uncorrected; parametric t-test), which did not survive multiple comparison correction.

|  | M Night | D Night | p value (uncorrected) |
| --- | --- | --- | --- |
| Sleep onset (min) | 11.89 ± 1.24 | 12.17 ± 1.22 | 0.825 |
| TST (min) | 57.41 ± 3.37 | 57.46 ± 3.29 | 0.992 |
| S1 (%) | 3.92 ± 0.49 | 6.61 ± 0.87 | 0.008 * |
| S2 (%) | 30.36 ± 3.55 | 29.23 ± 3.13 | 0.704 |
| S3 (%) | 23.65 ± 2.49 | 26.53 ± 2.60 | 0.110 |
| S4 (%) | 22.59 ± 3.50 | 17.42 ± 3.23 | 0.052 |
| SWS (%) | 46.24 ± 3.07 | 43.95 ± 3.11 | 0.422 |
| REM (%) | 0.41 ± 0.41 | 0 ± 0 | 0.328 |
| WASO (%) | 2.35 ± 0.46 | 3.74 ± 0.72 | 0.059 |
| Arousals (#) | 6.61 ± 1.57 | 5.70 ± 1.18 | 0.526 |
| SE | 80.94 ± 1.93 | 79.80 ± 1.82 | 0.491 |

## Discussion

We investigated the neural signatures associated with odor cueing of declarative memories during NREM sleep. To this goal, we compared electrophysiological features to cueing with an odor that was associated with training on a declarative memory task against cueing with an odor that was associated with a procedural random finger-tapping task designed to minimize learning. Odor cueing of declarative memories increased the rate of the sleep spindles over centro-parietal areas, as well as spindle duration over frontal areas. As for SOs, declarative memory cueing resulted in increased peak amplitudes. Furthermore, the coupling between SOs and spindles was enhanced, such that spindle increases were accumulated mainly around the SO up states compared to other phases of the SO cycle. Interestingly, all demonstrated changes were strongest in the first half of the cueing periods. In summary, using a well-controlled experimental design, we show that declarative memory-associated odor cues induce spatially congruent enhancements of spindle and SO activity, as well as of their temporal coupling. Characterizing these changes is a crucial step towards a better understanding of the neural underpinnings of both cued and spontaneous memory consolidation. Importantly, our experimental paradigm was designed to minimize potential confounding factors. Since the paradigm effectively controls for unspecific effects of presenting odors during sleep, our results are likely to genuinely reflect memory re-processing during sleep.

Since its initial demonstration^5^, the beneficial effect of memory cueing using odors has been shown for creativity tasks^29^, fear extinction^17^, declarative memory in a real-life setting^30, 31^. The beneficial effect of cueing on memory has been shown to be odor-specific^16, 32^, can be expressed unilaterally in the brain^15^, and is protected against interference when applied during sleep^6, 32–34^. While in the current study, the effects of declarative memory odor cueing were not expressed in significantly improved declarative memory performance (**Supplementary Figure 1**), this does not preclude the presence of robust electrophysiological signatures of the induced memory processes, as demonstrated in prior studies^13, 14^. However, electrophysiological evidence for the effects of odor cueing is sparse.

Concerning cueing effects on discrete events, two studies have shown an increase of spindle density in parietal^13^ and occipital areas^14^. The present study reveals changes in spindle rate over more widespread areas, as well as duration changes over frontal instead of only occipital areas. Such differences are not surprising, as sleep spindles are rather local events and their topology may change according to the learned content, properties of its presentation (e.g., location in the visual field), and details of the cueing. Despite our meticulous efforts to focus on fast spindles, it is important to note that we cannot guarantee that portions of slow spindles were captured by our event detection. The rather unexpected result of increased spindle duration over frontal regions (while spindle rate was increased centro-parietally) may thus reflect a change in slow rather than fast spindles. Concerning SOs, while we did not detect a steepening of SO slopes in response to declarative memory odor cueing, as has previously been shown by Rihm et al.^16^, we observed increased SO amplitudes, which might point to a similar phenomenon, as amplitude and slope of oscillations are intrinsically related. Interestingly, cueing-induced SO amplitude increases occurred over centro-parietal areas and overlapped with increases in spindle rate. Our follow-up analysis showed that SO amplitude and the likelihood of the occurrence of a coupled sleep spindle are indeed related. Together, these results suggest that the two phenomena are not independent, but that higher SO amplitudes drive increased spindle rates or vice versa.

Turning to the more fine-grained temporal coupling of SOs and spindles, cueing-evoked changes in this coupling have been shown in several auditory stimulation studies, where the cue itself is likely to elicit an SO and a nested spindle^19, 35^. Only one study so far has revealed changes in phase-amplitude coupling in response to odor cueing^15^. Our observation that the preferred phase of the spindle coupling is shifted towards the positive peak of the SO is in line with this study. Furthermore, spindles that occurred during SOs were increased in particular during their up states. How these phenomena are linked mechanistically is unclear and beyond the scope of our study. Interestingly, the strength of the coupling did not significantly change due to cueing. This might result from induced spindles being broadly distributed around the preferred phase.

Most cueing-induced changes in neural activity were strongest, and to large parts only detectable, during the first half of the stimulation period. A similar phenomenon has been described in a previous study in which odor cueing-induced increases in delta and fast spindle activity were more dominant in the first seconds after stimulation onset^16^. These observations may suggest that the cueing effect is relatively short-lived, possibly due to fast-acting habituation to the odor or refractory periods of the involved neural mechanisms (e.g., spindle initiation^36^). The fact that cueing-induced changes in the preferred coupling angle of spindles to SOs were only significant across the whole stimulation period might be explained by narrowing down the analyzed time windows resulting in low numbers of events and thus low statistical power. Nevertheless, the predominant clustering of induced changes to the first half of the cueing period suggests that shorter stimulation time windows may be more optimal for odor cueing. This would allow for a larger number of assessable trials and might even enhance the behavioral effect of the odor cueing.

In this study, we put particular emphasis on how we normalized and statistically compared our data between conditions. Previous studies have often used breaks between stimulations (sometimes called “odor-off” periods) to normalize responses to stimulation, and another (sham or vehicle-only) night as the control condition. In our strategy, vehicle time periods used for normalization are separated from the odor periods by a post-stimulation break of the same length. We believe this approach provides a cleaner comparison, since brain activity elicited by the odor could persist after odor offset. Our analysis strategy (normalizing against vehicle in the same night and contrasting against normalized data acquired using a control odor in another night) controls for differences between individuals, nights, odor familiarity, memory-unspecific effects of odors, and, because vehicle periods were interleaved with odor periods, also for time-of-night effects.

### Limitations and future research

One limitation of the current study is the intrinsic temporal vagueness of the olfactory cueing modality. The odor takes time to travel through the air pipes of the olfactometer and its perception is subject to the respiratory rhythm^37^. Improved ways to release odors with high temporal accuracy together with respiratory monitoring could provide greater consistency across stimulation trials. Another limitation is the lack of behavioral performance differences between the cueing conditions, for which there are various potential explanations. On the one hand, both odors may have indiscriminately been associated with broad contextual memories of the few hours before sleep. This way, despite not having been paired directly, Odor M may have inadvertently improved performance on the declarative memory task. Another explanation may be that testing for performance differences soon after the cueing intervention might have obscured effects on memory. Two recent studies have shown an expression of behavioral effects of auditory memory cueing only after several hours^35^ or even days^38^. In one of these studies, the category of the cued memory items could be decoded during cueing-evoked neural responses, while improvements in memory only emerged after another full night of sleep^35^. Finally, low statistical power common to most neuroscientific studies may hinder the detection of small effects. Nevertheless, the lack of a behavioral effect of cueing allows for the possibility that the observed electrophysiological changes are not directly related to memory reactivation. However, as our results are broadly in line with prior cueing-induced changes in studies that did show a behavioral effect ^15, 16^, we assign a low likelihood to this scenario.

In summary, we extend the sparse prior literature on the neural effects of odor cueing by analyzing various features of slow oscillations, sleep spindles, and their temporal coupling in response to stimulation. We demonstrate that cueing with a declarative memory-associated odor increases spindle rate and duration, SO amplitude, and the number of spindles occurring specifically around SO up states. We propose slow oscillation amplitude enhancements as well as linked increases in the rate of sleep spindles as a potential mechanism of cueing-induced memory benefits. In addition to advancing our knowledge of memory processing during sleep, we hope our results also pave the way for a more frequent use of odor cueing in electrophysiological investigations of memory consolidation.

## Acknowledgments

This research was financially supported by the ANID FONDECYT Regular grant 1171320, year 2017. Additionally, A.S-C. was financially supported by the scholarship BECAS DE DOCTORADO VRI – ESCUELA DE GRADUADOS, Pontificia Universidad Católica de Chile. D.M.B. was funded by ANID FONDECYT Postdoctorado grant 3200051, year 2020 and G.A.W. was supported in part by the Centro Basal FB0008. J.G.K., S.H., and J.B. were supported by grants from the Deutsche Forschungsgemeinschaft (German Research Council, Tr-SFB 654 “Sleep and Plasticity”).

