## Supplementary for "Odor cueing of declarative memories during sleep enhances coordinated spindles and slow oscillations"

### Supplemental material

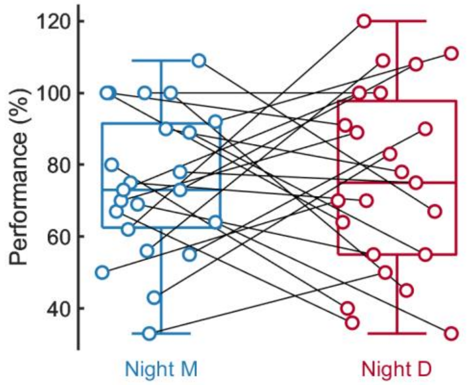


***Supplementary Figure 1: Declarative memory performance.*** *Box plots show median and 25th and 75th percentiles. Circles represent the individual performance on the memory task, calculated as the number of correctly answered card pairs during post-sleep testing (Recall) relative to immediate recall after the learning session (Learning). Lines connect the subject's performances between M and D night.*

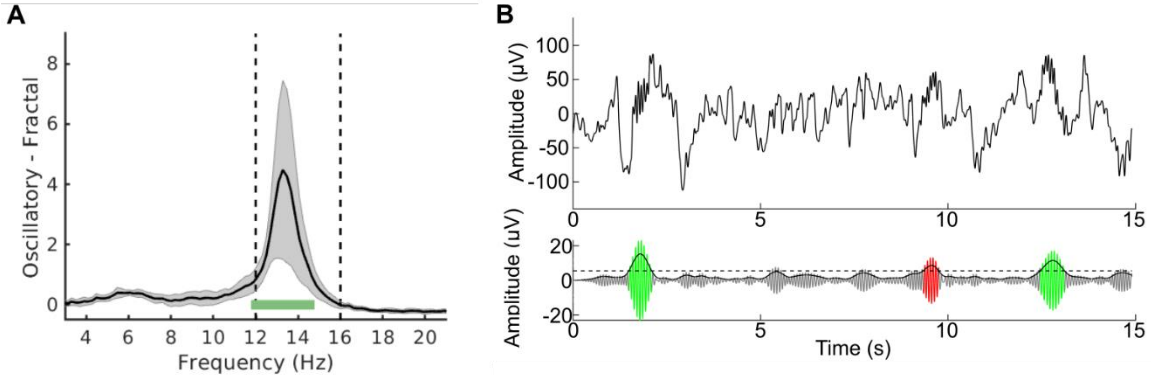


***Supplementary Figure 2: Example of subject-specific determination of spindle peak frequency.*** *For each subject, peak frequencies were identified by irregular-resampling auto-spectral analysis (IRASA) of data recorded from an a-priori defined central cluster of electrodes. (**A**) Oscillatory power component (mean ± SD) across these electrodes. The spindle peak was defined as the maximum power within the canonical fast spindle range (12 - 16 Hz (vertical dashed lines). A 3-Hz frequency band (green horizontal bar) centered around the defined peak served as a bandpass for spindle event detection. (**B**) A segment of data from the central cluster (top) and result of the spindle detection (bottom). The horizontal dashed line represents the amplitude threshold for spindle candidates to cross. Green and red overlays respectively show the signals of valid and invalid spindle candidates.*

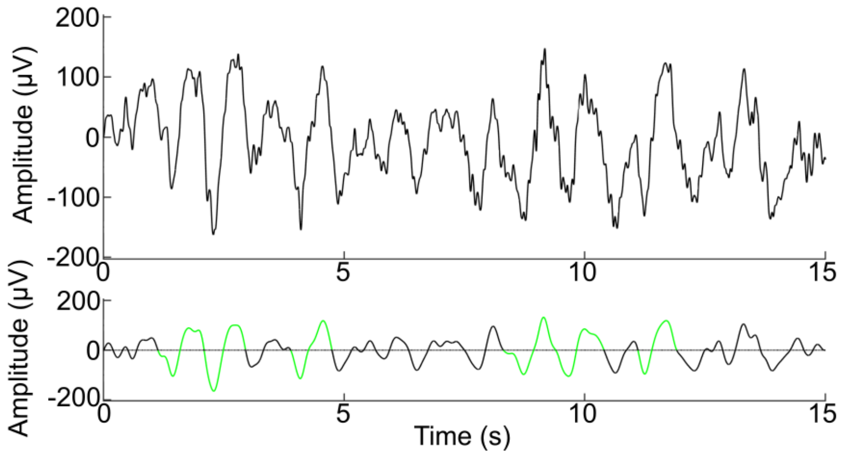


***Supplementary Figure 3: Slow oscillation event detection.*** *Example of wideband data recorded at electrode Fz (0.1 - 45 Hz; upper panel) and the same data filtered in the SW band (0.5 - 4 Hz, lower panel). Individual slow oscillations (green) were detected on SW-filtered signals as described in the methods.*

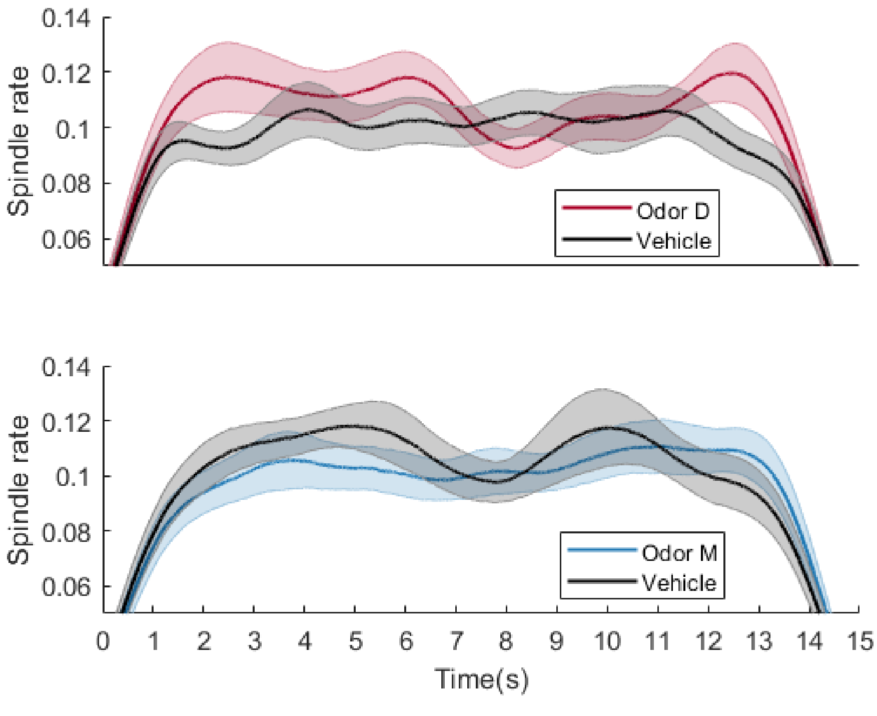


***Supplementary Figure 4: Average spindle rate over time.*** *Time series of the trial average spindle rate over the defined central cluster of electrodes. Each spindle occurrence is considered as a spike (taking the analogy of a firing neuron) and convoluted with a Gaussian kernel for smoothing. The upper panel shows the average spindle rate in D Night where Odor D (red) and the vehicle (black) were presented. The bottom panel shows the average spindle rate in M Night where Odor M (blue) and vehicle (black) were presented. Spindles starting before cero or ending after 15 s were*

*not considered. For D Night, an increase in the spindle rate is observed for odor D in the first half of the trial. After 7 to 12 seconds, a decrease is presented probably related to a refractoriness of the sleep spindles.*

***Supplementary Table 1: Number of slow oscillation events per trial for each subject***

| Subject | Mean events per trial Odor D Night 0-7.5 s | Mean events per trial Vehicle D Night 0-7.5 s | Mean events per trial Odor M Night 0-7.5 s | Mean events per trial Vehicle M Night 0-7.5 s | Mean events per trial Odor D Night 0-15 s | Mean events per trial Vehicle D Night 0-15 s | Mean events per trial Odor M Night 0-15 s | Mean events per trial Vehicle M Night 0-15 s |
|---|---|---|---|---|---|---|---|---|
| **1** | 0.73 | 0.92 | 0.87 | 1.07 | 1.49 | 2.06 | 1.79 | 1.93 |
| **2** | 0.72 | 0.58 | 0.54 | 0.58 | 1.28 | 1.37 | 1.23 | 1.36 |
| **3** | 1.22 | 0.67 | 0.72 | 0.68 | 1.95 | 1.25 | 1.46 | 1.73 |
| **4** | 0.55 | 1.08 | 0.68 | 0.68 | 0.91 | 2.19 | 1.40 | 1.21 |
| **5** | 0.37 | 0.43 | 0.74 | 0.66 | 1.35 | 1.24 | 1.73 | 1.80 |
| **6** | 0.56 | 0.44 | 0.37 | 0.31 | 0.98 | 0.93 | 0.78 | 0.68 |
| **7** | 1.21 | 0.77 | 0.66 | 1.08 | 1.91 | 1.74 | 1.69 | 1.93 |
| **8** | 0.89 | 0.72 | 0.55 | 0.68 | 1.76 | 1.44 | 1.37 | 1.62 |
| **9** | 0.57 | 0.76 | 0.68 | 0.44 | 1.22 | 1.50 | 1.18 | 1.23 |
| **10** | 0.72 | 0.85 | 0.57 | 0.80 | 1.55 | 1.66 | 1.20 | 1.56 |
| **11** | 0.94 | 0.57 | 0.58 | 0.77 | 1.50 | 1.19 | 1.19 | 1.58 |
| **12** | 0.89 | 0.96 | 0.89 | 0.73 | 1.94 | 2.04 | 1.91 | 1.62 |
| **13** | 0.74 | 0.40 | 0.81 | 0.66 | 1.57 | 1.16 | 1.68 | 1.27 |
| **14** | 0.55 | 0.53 | 0.65 | 0.59 | 1.44 | 1.30 | 1.52 | 1.37 |
| **15** | 0.72 | 0.69 | 0.73 | 0.71 | 1.61 | 1.39 | 1.74 | 1.48 |
| **16** | 0.93 | 0.71 | 0.88 | 0.64 | 1.52 | 1.84 | 1.74 | 1.03 |
| **17** | 0.78 | 1.04 | 0.80 | 0.79 | 1.64 | 1.84 | 1.72 | 1.60 |
| **18** | 0.48 | 0.51 | 0.68 | 0.93 | 0.86 | 1.53 | 1.93 | 1.86 |
| **19** | 0.66 | 0.84 | 0.79 | 0.52 | 1.55 | 1.67 | 1.66 | 1.45 |
| **20** | 0.79 | 0.59 | 0.88 | 0.90 | 1.96 | 1.52 | 1.70 | 1.76 |
| **21** | 0.61 | 0.54 | 0.64 | 0.46 | 1.27 | 1.16 | 1.43 | 0.98 |
| **22** | 0.83 | 0.99 | 0.98 | 0.71 | 1.78 | 1.77 | 2.08 | 1.45 |
| **23** | 0.69 | 0.59 | 0.63 | 0.61 | 1.37 | 1.32 | 1.18 | 1.11 |
| **Mean** | 0.75 | 0.70 | 0.71 | 0.70 | 1.50 | 1.53 | 1.54 | 1.46 |
| **Std** | 0.0088 | 0.0085 | 0.0060 | 0.0079 | 0.0134 | 0.0140 | 0.0131 | 0.0137 |

*The first four columns show the mean SO counts per trial for the first half of the stimulation period (on which we focused for large parts of the analysis; values are provided separately for Odor D and M nights, as well as Odor and Vehicle stimulation). The last four columns show the same metrics for the entire trial. Values represent the mean across all the channels of an anatomically defined central cluster (see Figure 2A). Analyses showed no significant main effects or interactions (all $p > 0.25$; repeated-measures ANOVAs on half or full trial data, incorporating the two within-subject factors Odor D/M and Odor/Vehicle).*

***Supplementary Table 2: Number of spindle events per trial for each subject.***

| Subject | Mean events per trial Odor D Night 0-7.5 s | Mean events per trial Vehicle D Night 0-7.5 s | Mean events per trial Odor M Night 0-7.5 s | Mean events per trial Vehicle M Night 0-7.5 s | Mean events per trial Odor D Night 0-15 s | Mean events per trial Vehicle D Night 0-15 s | Mean events per trial Odor M Night 0-15 s | Mean events per trial Vehicle M Night 0-15 s |
|---|---|---|---|---|---|---|---|---|
| **1** | 0.23 | 0.37 | 0.43 | 0.22 | 0.59 | 0.84 | 0.73 | 0.66 |
| **2** | 0.56 | 0.37 | 0.50 | 0.36 | 0.90 | 1.05 | 1.13 | 0.87 |
| **3** | 0.33 | 0.31 | 0.40 | 0.45 | 0.73 | 0.80 | 0.77 | 0.83 |
| **4** | 0.08 | 0.33 | 0.18 | 0.43 | 0.35 | 0.58 | 0.54 | 0.69 |
| **5** | 0.36 | 0.23 | 0.22 | 0.29 | 0.78 | 0.65 | 0.64 | 0.78 |
| **6** | 0.27 | 0.37 | 0.20 | 0.36 | 0.62 | 0.80 | 0.66 | 0.56 |
| **7** | 0.49 | 0.44 | 0.52 | 0.44 | 0.94 | 0.82 | 0.88 | 0.86 |
| **8** | 0.39 | 0.34 | 0.30 | 0.27 | 0.65 | 0.67 | 0.67 | 0.59 |
| **9** | 0.44 | 0.43 | 0.34 | 0.60 | 0.93 | 0.87 | 0.87 | 1.11 |
| **10** | 0.48 | 0.42 | 0.34 | 0.29 | 0.78 | 0.86 | 0.59 | 0.70 |
| **11** | 0.49 | 0.31 | 0.31 | 0.27 | 0.82 | 0.51 | 0.64 | 0.69 |
| **12** | 0.32 | 0.48 | 0.32 | 0.40 | 0.62 | 0.68 | 0.70 | 0.62 |
| **13** | 0.64 | 0.28 | 0.24 | 0.45 | 1.11 | 0.63 | 0.65 | 1.09 |
| **14** | 0.53 | 0.27 | 0.33 | 0.39 | 0.99 | 0.83 | 0.68 | 0.82 |
| **15** | 0.50 | 0.56 | 0.44 | 0.46 | 1.05 | 1.04 | 0.99 | 0.88 |
| **16** | 0.49 | 0.28 | 0.40 | 0.44 | 0.90 | 0.66 | 0.69 | 0.56 |
| **17** | 0.45 | 0.38 | 0.35 | 0.32 | 0.85 | 0.73 | 0.68 | 0.62 |
| **18** | 0.27 | 0.50 | 0.45 | 0.50 | 0.63 | 0.91 | 0.94 | 1.07 |
| **19** | 0.63 | 0.39 | 0.46 | 0.40 | 1.20 | 0.81 | 1.06 | 0.81 |
| **20** | 0.40 | 0.41 | 0.44 | 0.46 | 0.80 | 0.79 | 0.72 | 0.79 |
| **21** | 0.43 | 0.30 | 0.29 | 0.55 | 0.82 | 0.65 | 0.73 | 0.92 |
| **22** | 0.36 | 0.42 | 0.33 | 0.31 | 0.73 | 0.82 | 0.71 | 0.85 |
| **23** | 0.23 | 0.32 | 0.17 | 0.36 | 0.52 | 0.42 | 0.42 | 0.64 |
| **Mean** | 0.41 | 0.37 | 0.35 | 0.39 | 0.80 | 0.76 | 0.74 | 0.78 |
| **SEM** | 0.0058 | 0.0035 | 0.0043 | 0.0040 | 0.0084 | 0.0065 | 0.0071 | 0.0069 |

*The first four columns show the mean spindle counts per trial for the first half of the stimulation period (on which we focused for large parts of the analysis; values are provided separately for Odor D and M nights, as well as Odor and Vehicle stimulation). The last four columns show the same metrics for the entire trial. Values represent the mean across all the channels of an anatomically defined central cluster (see Figure 2A). Analysis showed a borderline significant effect of declarative memory cueing on spindle counts ($p = 0.053$, $\eta^2 = 0.064$; interaction Odor D/M × Odor/Vehicle; repeated-measures ANOVA performed on data from the first half of the stimulation period). No other comparisons were significant (all $p > 0.19$).*
